# Genetic influences on brain activation and large-scale functional connectivity during nociceptive processing: a twin study

**DOI:** 10.1101/2021.01.08.425878

**Authors:** Gránit Kastrati, Jörgen Rosén, William H. Thompson, Xu Chen, Henrik Larsson, Thomas E. Nichols, Irene Tracey, Peter Fransson, Fredrik Åhs, Karin B. Jensen

## Abstract

Nociceptive processing in the human brain is a signal that enables harm avoidance, with large interindividual variance. The relative contributions of genes and environment to the neural structures that support nociception have not been studied in twins previously. Here, we employed a classic twin-design to determine brain structures influenced by additive genetics. We found genetic influences on nociceptive processing in the midcingulate cortex, bilateral posterior insulae and thalamus. In addition to brain activations, we found genetic contributions to large-scale functional connectivity during nociceptive processing. We conclude that additive genetics influence specific aspects of nociceptive processing, which improves our understanding of human nociceptive processing.

## Introduction

Nociceptive processing is important for survival as it provides an organism with information about potential or actual tissue damage. The neural processes underlying this capacity are evolutionary conserved, as evolved nociceptive systems are observed in a variety of species (Walters & Williams, 2019). In humans, neuroimaging studies have established a large network of brain regions that consistently activate in response to nociceptive information (Jensen et al., 2016). Most such activations are evoked independently of type of nociceptive input and can be found in infants with minimal prior exposure to pain (Goksan et al., 2015). This suggests that genes modulate basic aspects of nociceptive processing in the human brain.

There is considerable variation in nociception between individuals and attempts have been made to determine the genetic influence on such differences (Mogil, 2012). The genetic influence on sensitivity to experimental pain, for example, has been investigated by comparing identical and fraternal twins and estimated to 26% for heat- and 60% for cold-induced pain (Nielsen et al., 2008). Another study found similar genetic influence on individual sensitivity to pain, ranging from 22-55% depending on pain modality (Norbury, MacGregor, Urwin, Spector, & McMahon, 2007). Studies that link single-nucleotide polymorphisms to functional neuroimaging data e.g. (Oertel et al., 2008; Vachon-Presseau et al., 2016; Zubieta et al., 2003) and studies of rare genetic mutations that affect pain perception (Salomons, Iannetti, Liang, & Wood, 2016) suggest that our genes influence nociceptive processing, and our subjective experience of pain. Yet, the specific neural mechanisms and magnitude of such influence needs to be determined.

The experience of pain likely involves cross-communication between both nociceptive and non-nociceptive brain regions (Geuter et al., 2020; Kucyi & Davis, 2015). To capture genetic influences on nociceptive processing, it is therefore relevant to move beyond mere activations in specific brain regions, to also consider their interactions. Recent advances in the neurosciences have seen a rapid increase in studies that model the brain as a large-scale network, which allows for estimating the degree of the interaction or cross-communication between brain regions and/or sub-networks (Bullmore & Bassett, 2011; Sporns, 2013). For example, the default-mode network that consistently activates during rest and deactivates when engaged in a task, show increased deactivation during painful tasks (Kong et al., 2010; Kucyi, Salomons, & Davis, 2013). Recent findings also show decreased functional connectivity between the primary somatosensory cortex and the default-mode network in chronic low back pain (Kim et al., 2019). Several studies have elucidated the relationship between genetics and functional brain network topology by means of functional Magnetic Resonance Imaging (fMRI), both for resting-state (Fornito et al., 2011; Glahn et al., 2010; Miranda-Dominguez et al., 2018; Reineberg, Hatoum, Hewitt, Banich, & Friedman, 2019; Xu et al., 2017) and experimental tasks (Alstott, Breakspear, Hagmann, Cammoun, & Sporns, 2009; Colclough et al., 2017; Yang et al., 2016). Estimates of the genetic influence of resting-state brain networks are replicable across studies and imply genetic influences on large-scale networks (Adhikari et al., 2018).

In this study, we estimated the genetic influence on nociceptive processing in the brain. A total of 246 twins (56 identical pairs; 67 fraternal pairs) participated in a fMRI study that included an aversive conditioning paradigm using electrical shocks. The aim was to estimate the genetic influence on 1) neural responses in pain processing regions and 2) whole-brain functional connectivity during nociceptive processing, as described in our preregistration protocol (https://osf.io/zesw5). To achieve our first aim, we constrained our analysis to pain processing regions defined independently of the current study (Wager et al., 2013). Regarding the second aim, we used a whole-brain parcellation scheme to study pain-evoked functional connectivity.

## Materials and Methods

### Subjects

Twins between ages 20 and 60 years were recruited for the present study through the Swedish Twin Registry (STR). The STR contains more than 194 000 twins and represents an epidemiological resource for the study of genetic and environmental influences on human traits, behaviors, and diseases (https://ki.se/en/research/the-swedish-twin-registry). Twin pairs with known zygosity were selected based on their capability to undergo magnetic resonance imaging as well as screened for substance abuse, ongoing psychological treatment or medicine affecting emotion or cognition. Only same-sex twin pairs were included in this study and after initial screening, 305 participants were recruited to the study and underwent fMRI scanning. Imaging data were excluded from the analysis if one of the following criteria were fulfilled (i) excessive amount of head motion (more than 50 % of the data frames contained framewise displacement above 0.5 mm) (n=16). (ii) presence of outliers in terms of amplitude of brain responses. We here used the median absolute deviation method (Leys, Ley, Klein, Bernard, & Licata, 2013) to detect and outliers (here meaning a mean blood-oxygenated-level-dependent (BOLD) response deviating more than 3 times the medial standard deviation across the whole sample). Imaging data from participants deemed to be outlies were removed together with data from their co-twin and not used in the subsequent analysis (n=8). (iii) missing data / incomplete data collection from both twin pairs (n=35). The final sample (n=246) included 56 identical (35 female, 21 male) twin pairs (age: M=34, SD=8) and 67 fraternal (39 female, 28 male) twin pairs (age: M=33, SD=11). All participants provided written informed consent in accordance with the Uppsala Ethical Review Board Guidelines. Participants received reimbursement of SEK 1000 (roughly equal to 100 USD) for their participation.

### Brain imaging

Imaging data were acquired using a 3.0 T scanner (Discovery MR750, GE Healthcare) and an 8-channel head-coil. Foam wedges, earplugs and headphones were used to reduce head motion and scanner noise. We acquired T1-weighted structural images with whole-head coverage, TR=2.4s, TE=2.8s, acquisition time 6.04 min and flip angle 11 (degrees). Functional images were acquired using gradient echo-planar-imaging (EPI), TR = 2.4 s, TE = 28ms, flip angle = 80 (degrees), with 47 seven volumes acquired with slice thickness 3.0 mm^3^ (no spacing, axial orientation, phase-encoding direction A/P). The slices were acquired in an interleaved ascending order. Higher order shimming was performed, and five dummy scans were acquired before the experiment.

### Stimuli and Contexts

Visual stimuli were presented on a flat screen in the MR scanner via a projector (Epson EX5260) (Fig. S1). The computer running the stimulus presentation used a custom version of Unity (version 5.2.3, Unity Technologies, San Francisco, CA) and communicated with BIOPAC for electrical stimuli (BIOPAC Systems, Goleta, CA) through a parallel port interface. The software for the parallel port interface was custom made and used standard .NET serial communication libraries by Microsoft (Microsoft Corporation, Albuquerque, New Mexico).

### fMRI paradigm design

Noxious electrical stimuli were administered as part of a fear conditioning procedure. The paradigm was used to test genetic aspects of fear acquisition and results that focus on neural responses to trials that did not include an electrical shock will be reported elsewhere. Two virtual characters served as visual stimuli (CS) and were presented at a distance of 2.7 m projected on a screen in the MR scanner (Fig. S1). One of the virtual characters served as the aversive cue (CS+) and preceded the electrical stimuli whereas the other virtual character served as a safety cue (CS-). Stimuli serving as CS+ and CS-was counterbalanced across participants. Each of the cues appeared for 6s. Participants were not told which character would be associated with electrical shocks. Prior to the conditioning phase, a habituation phase took place, during which each CS was presented four times without any electrical shocks. During conditioning, each cue type was displayed 16 times. Eight of the aversive cues co-terminated with presentation of the electrical shock (US) and eight of the aversive cues did not include a shock. Four stimulus presentation orders were used to counterbalance the timing of CSs across subjects. An inter-stimulus interval (randomized jittering) followed each trial, with no cues present for 8-12s. Total duration for the conditioning task was 9 minutes and 47 seconds. The initial 8 presentations (habituation) were not considered for this analysis.

The electrical shocks were delivered to the distal part of the participant’s left volar forearm (adjacent to the wrist) via radio-translucent disposable dry electrodes (EL509, BIOPAC Systems, Goleta, CA). As the present study also served to investigate fear acquisition, i.e., neural responses to trials that did not include electrical stimulation) (to be published elsewhere), the US presentation was brief (16 ms). Shock delivery was controlled using the STM100C module connected to the STM200 constant current stimulator (BIOPAC Systems, Goleta, CA), using a unipolar pulse with a fixed duration of 67 Hz. The physical voltage was individually calibrated before the experimental task using an ascending staircase of electrical currents until shocks were rated as ‘aversive’ (Rosen, Kastrati, Reppling, Bergkvist, & Ahs, 2019). After finding the physical voltage that participants rated as aversive, this parameter was kept constant throughout the experiment. The determined average electrical voltage was M=31V, SD=7 across participants.

### Analysis of fMRI imaging data

Analyses of fMRI-data were performed using SPM12 (Welcome Department of Cognitive Neurology, University College, London, https://www.fil.ion.ucl.ac.uk/spm). Preprocessing of functional image volumes included interleaved slice time correction, realignment, co-registration to the T1-weighted image, spatial normalization to Montreal Neurological Institute (MNI) space (MNI152NLin6Asym), and spatially smoothed with an 8mm Gaussian kernel.

In the first-level analysis, an event-related approach was used to estimate BOLD responses during nociceptive processing. Three event types were modeled, using separate regressors: the aversive cue that preceded the US (CS+_US_), the same CS+ that did not precede the US (CS+_no US_), and the electrical shock itself (US). Note that the aversive cue (CS+) co-terminated with the onset of the US 50% of the times. The duration of the visual cue (CS+) was set to 6 seconds and the US to 3 seconds. The first-level contrast for each participant that was latter used to estimate the genetic influence *h*^*2*^ on nociceptive processing per se was modeled as (CS+_US_ & US > CS+_no US_). Since the aversive cue (CS+_US_) was immediately followed by the US, without any delay, the CS+_US_ and US were combined. The same visual cue (CS+_no US_), not followed by the US, was then subtracted in order to estimate the neural correlates to nociceptive processing *per se*. The group-level result for the same contrast is found in Table S1 and Fig. S2. The statistical significance threshold was set to *P* < 0.05, family-wise error corrected (FWE) for multiple comparisons. Anatomical labeling of significantly activated brain regions were performed using the SPM Anatomy toolbox v.2.2c (Tzourio-Mazoyer et al., 2002).

### Defining the functional connectome in response to nociceptive input

To investigate task-specific functional connectivity, the CONN functional connectivity toolbox was used (Whitfield-Gabrieli & Nieto-Castanon, 2012) (http://www.nitrc.org/projects/conn, version 18b). As input to the CONN toolbox, we used the same preprocessing pipeline as outlined above except for removing the spatial smoothing. This decision was to minimize a spurious increase in local connectivity that would be induced otherwise. Subsequently, image data underwent ART-based outlier detection of volumes (version 2015-10) followed by image scrubbing. For the scrubbing procedure, we used a liberal threshold of the 99^th^ percentile of normative sample, with a global-signal *z*-value threshold of 9 standard deviations and a subject motion threshold of 2mm. Next, confounders were removed from the data. These consisted of the effect of each task (in order to remove constant task-induced responses in the BOLD signal), cerebrospinal fluid, white matter, SPM covariates (6 motion parameters and their quadratic effect) and regressors for scrubbing per individual (one regressor for each volume deemed a potential outlier; from zero to a maximum of 25 regressors per individual). Finally, image data was low-pass filtered [0.008, 0.09]. BOLD time-series were extracted using a parcellation scheme with 400 nodes (Schaefer et al., 2018). We computed first-level weighted ROI-to-ROI functional connectivity (wFC) by computing task-specific bivariate correlation using weighted Least Squares (WLS), with weights defined as condition timeseries convolved with a canonical hemodynamic response function. Results were Fisher-transformed correlation coefficients between each pair of nodes. The first-level contrasts were modelled in the same way as described above for brain activations (CS+_US_ & US > CS+_no US_). Fig. 2A and 2C shows the group-level result for the same contrast. For visualization purpose, we computed the within-network and between-network sum of functional connectivity between each pair of networks (Fig. 2C). For each network, say *A* and *B*, we sum the functional connectivity between *A* and *B* and divide by the number of nodes contained in the two networks. If *A=B*, the result is the sum of the within-network connectivity; otherwise, the result is the between-network connectivity.

**Fig 1.**
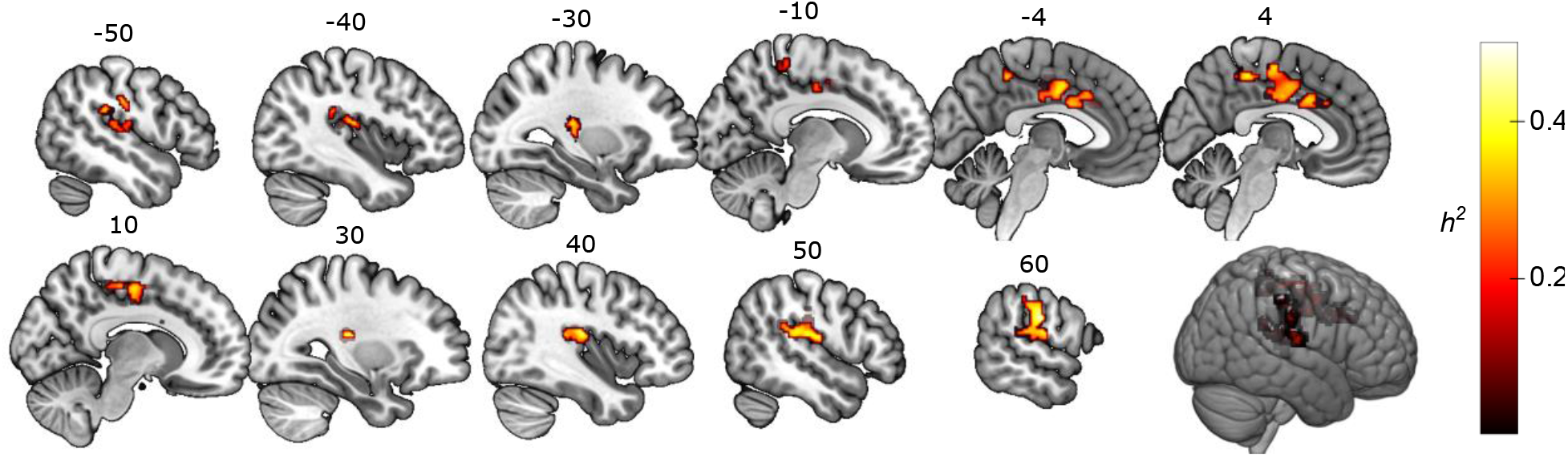
Twin-data brain regions with genetic influences during nociceptive processing. Sagittal view of clusters with significant genetic influence, including the contralateral somatosensory cortex, bilateral dorsal posterior insulae, anterior and midcingulate cortex. The threshold was set at p<.05, FWE-corrected for multiple comparisons at the cluster-level. The heat bar represents *h*^2^ heritability values.

**Figure 2.**
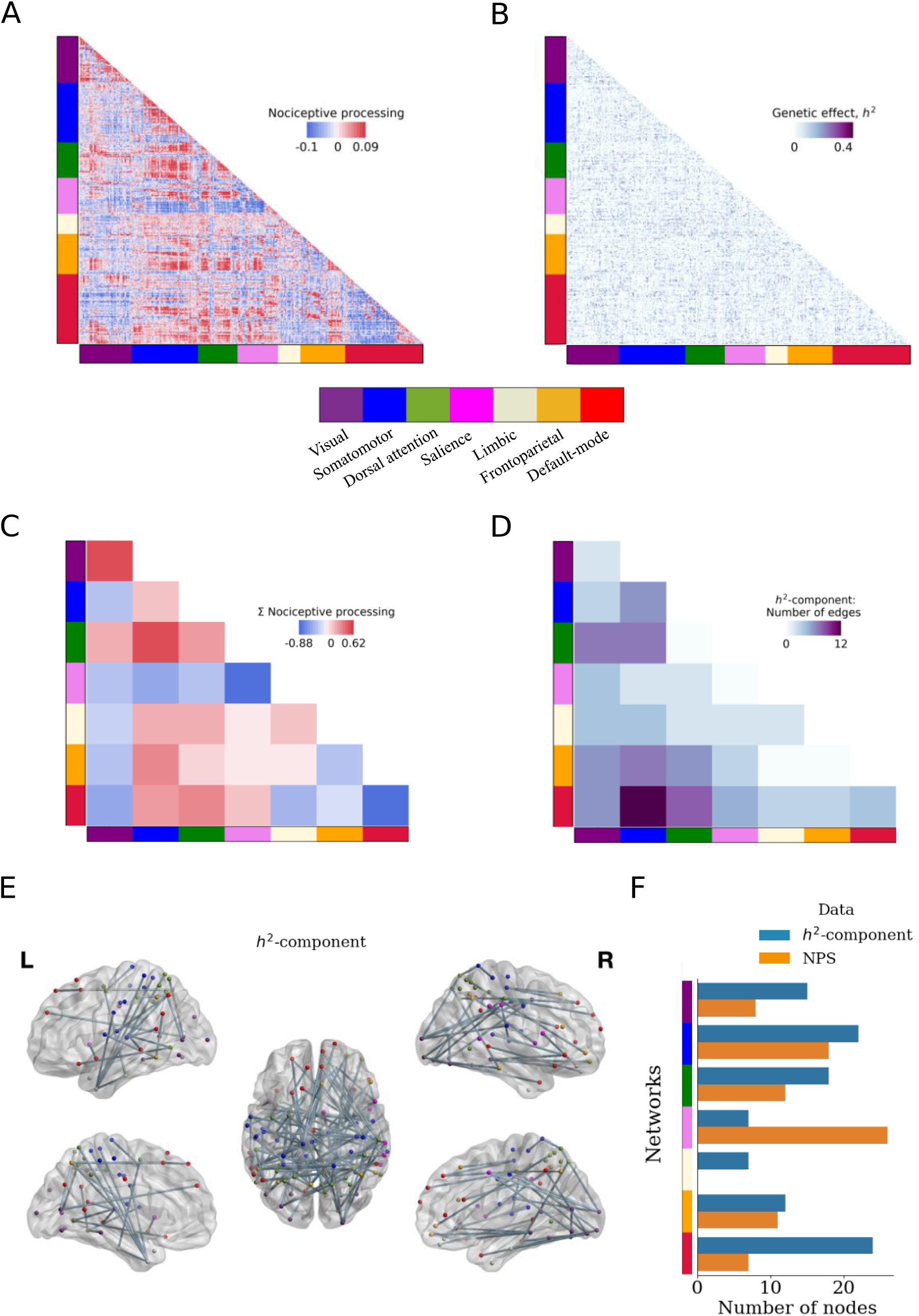
Twin-data functional connectivity during nociceptive processing. **(A)** Group-averaged functional connectivity (FC) during nociceptive processing. Positive values (red) indicate edges with stronger FC during nociceptive processing. **(B)** Unthresholded genetic influence (*h*^*2*^) for every edge in the functional connectivity during nociceptive processing. (**C**) Graphical summary of the functional connectivity results in (A). The diagonal squares represent the within-network and off-diagonal squares represent the between-network sum of functional connectivity during nociceptive processing. Positive values are represented by warm colors. Minimum and maximum values denote the mean +/- two standard deviations. (**D**) The number of edges in the connectivity cluster defined by genetic influence, called the *h*^*2*^-component, within and between networks. Dark color denotes higher number of edges. The largest number of edges was found between the somatomotor and default-mode network. **(E)** Brain graph representing the *h*^*2*^-component from (D). The edges comprise an *h*^*2*^-component that represents significant genetic influences on nociceptive processing (p<0.05, corrected, *h*^*2*^ threshold = 0.328). Nodes are color-coded according to the network definitions given in (Yeo et al., 2011). (**F**) The number of nodes in the parcellation scheme that overlap with the *h*^*2*^-component (blue) (defined with a threshold of *h*^*2*^ = 0.328) or the Neurologic Pain Signature (orange).

### Estimation of genetic influences on brain function

#### Exclusion of outliers

We identified univariate outliers in our data sample using the median absolute deviation method (Leys et al., 2013). Any participant with a mean BOLD response deviating more than 3 times the median standard deviation was removed as well as their respective twin (number of participants removed = 8). Included in the final analysis was a sample of 56 monozygotic (35 females, 21 males) and 67 dizygotic (39 females, 28 males) twin pairs.

In brief, the phenotypic variance can be decomposed into additive genetic variance (*A*) as genetic effects for a phenotype or trait that add up linearly, common or shared environmental variance (*C*) and unique environmental, or error variance (*E*) (Falconer & Mackay, 1996). Using the simplest Falconer’s formula, the A, C, and E-factors can be estimated by contrasting monozygotic-twin pair correlations with dizygotic-twin pair correlations. The A-factor can be identified because monozygotic-twins are genetically identical while dizygotic-twins share 50% of their co-segregating alleles on average. Additionally, we assume that a shared environmental contribution (C) is equally shared within pairs regardless if they are monozygotic or dizygotic twins. Finally, any variance not attributable to factors shared between twins (A and C), i.e., that make twins in pairs dissimilar, in the model assigned to the E-factor. The genetic influence (*h*^*2*^), is the proportion of a phenotypic variance explained by additive genetic effects, i.e. *h*^*2*^ is equal to A/(A+C+E). In the present study, we computed heritability using the APACE software package (Accelerated Permutation Inference for the ACE model) (Chen et al., 2019). APACE uses a non-iterative linear regression-based method based on squared twin-pair differences, with permutation-based multiple testing correction to control the family-wise error rate. For the mass-univariate analysis, for each first-level contrast described above, we used the Neurologic Pain Signature as *a priori* template for regions in which to test for significant differences in genetic influences between twin groups (Wager et al., 2013). The number of permutations was set to 1000 and we used the cluster-based inference in the APACE (Accelerated Permutation Inference for ACE models) software package (Chen et al., 2019) with cluster-forming threshold set to *p* < 0.05 based on the parametric likelihood ratio null-distribution. We additionally computed an estimate of the genetic influence of choice of threshold for the electrical stimulation using the mets package (Holst, Scheike, & Hjelmborg, 2016; Scheike, Holst, & Hjelmborg, 2014) implemented in R (R Core Team, 2017).

### Estimating the genetic effect on the functional connectome

All individual-level functional connectivity matrices (CS+_US_ & US > CS+_no US_) were entered into APACE (Chen et al., 2019) and the genetic influences was computed by fitting the model to each edge in the matrices. This resulted in a 400 by 400 symmetric matrix with *h*^*2*^ estimated for each edge. Subsequently, we used a method based on network-based statistics (Zalesky, Fornito, & Bullmore, 2010) to compute a significant cluster or ‘largest connected component’ of the *h*^*2*^ matrix. We ran 1000 iterations and re-computed the 400×400 *h*^*2*^ matrix with permuted twin identity. Finally, we computed the largest connected component of our observed *h*^*2*^ matrix and compared to the distribution of randomly generated *h*^*2*^ matrices, determining significance at α = 0.05. Of note, the network-based statistics approach requires a choice of a threshold for which below all values are set to zero and all values above are set to one. The usage of thresholds that are set too conservatively typically results in network components that are too small to be deemed significant compared to random networks. On the other hand, thresholds that are set too low results in very large network components that are biologically unrealistic. We found that the largest component broke at *h*^*2*^ = 0.328, however we show that there are larger components that are significant by computing components over several thresholds from *h*^*2*^ = 0.25 up to 0.32 in steps of 0.01 (see Fig. S5). For interpretability, we chose the component from the largest threshold, denoted the *h*^*2*^- component (*h*^*2*^ = 0.328) for visualization. To further aid interpretability, we computed the sum of within-network and between-network edges in the *h*^*2*^-component (Fig. 2D). All brain graphs where visualized using BrainNet Viewer (Xia, Wang, & He, 2013). Node labeling was done with the automated anatomical labeling (AAL) (Tzourio-Mazoyer et al., 2002) by taking the coordinates from the Schaefer parcellation (Schaefer et al., 2018) that overlap between the AAL and the *h*^*2*^- component.

### Notes on the preregistration

The aim of the current study as stated in the preregistration (https://osf.io/zesw5) was to characterize the genetic influence on functional connectivity in pain related brain regions. Our first approach was to use the automated online meta-analysis tool Neurosynth (Yarkoni, Poldrack, Nichols, Van Essen, & Wager, 2011) to determine the brain regions of interest. We here instead decided to use the Neurologic Pain Signature (Wager et al., 2013), since it is more well-defined and validated. In addition, instead of focusing the functional connectivity between brain regions related to pain, we took a whole-brain approach. This way, we could estimate the genetic influence on functional interactions between nociceptive and non-nociceptive brain regions. We decided furthermore to us weighted functional connectivity instead of generalized psychophysiological interactions (McLaren, Ries, Xu, & Johnson, 2012) since the former is conceptually simpler and sufficed for the present purpose. Finally, the permutation test based on network-based statistics (Zalesky et al., 2010) was added later, since element-wise (per edge) estimates of genetic influence assumes independence between edges, and would also match the cluster-based statistics from the univariate analysis.

## Results

### Genetic influence on brain activations during nociceptive processing

In response to nociceptive stimuli, we detected local increases in blood-oxygenated level-dependent (BOLD) fMRI signals in the bilateral anterior insulae, bilateral posterior insulae, cingulate cortex, thalamus, cerebellum, and the right amygdala (p<0.05, *FWE* corrected). For a full representation of all regions activated during nociceptive processing see *SI Appendix*, Table S1 and *SI Appendix*, Fig. S2. Estimates of the genetic influence on brain responses during nociceptive processing was constrained to brain regions defined by the Neurologic Pain Signature (Wager et al., 2013). Using permutation tests to assess the degree of genetic influence (*h*^2^ ranging from 0 to 1) on brain activation patterns (Chen et al., 2019), we found significant effects in the right (contralateral) postcentral gyrus (*h*^*2*^ = 0.52), right posterior insulae (*h*^*2*^ = 0.50), right superior temporal gyrus (*h*^*2*^ = 0.45), right supramarginal gyrus (*h*^*2*^ = 0.44), left postcentral gyrus (*h*^*2*^ = 0.54), left supramarginal gyrus (*h*^*2*^ = 0.52), left posterior insulae (*h*^*2*^ = 0.43), left superior temporal gyrus (*h*^*2*^ = 0.43), left anterior cingulate cortex (*h*^*2*^ = 0.46), right posterior-medial frontal gyrus (*h*^*2*^ = 0.41) and bilateral midcingulate cortex (*h*^*2*^ = 0.40) (Fig. 1) (see *SI Appendix*, SFig. 3 for an unthresholded image of the genetic influence, and *SI Appendix*, SFig. 4 for twin-pair correlations).

### Genetic influence on functional connectivity during nociceptive processing

During nociceptive processing we observed increases in functional connectivity *between* several brain networks, including the somatomotor and dorsal attention networks (Fig. 2A, C). The functional connectivity *within* the default-mode-network decreased during nociceptive processing and increased *within* the visual network. To estimate the genetic influence on functional connectivity, we used a permutation test based on network-based statistics (Zalesky et al., 2010). This approach allowed us to identify a cluster of connections from the full *h*^*2*^-matrix (Fig. 2B), where each connection represents the genetic influence on functional connectivity (Fig. 2D-F) (thresholded at p<0.05, corrected using 1000 permutations). The most conservative threshold where a significant cluster of connections could be determined (*h*^*2*^-component) was *h*^*2*^ = 0.328 (see *SI Appendix*, SFig. S5 for other thresholds). The edges of the *h*^*2*^-component linked together brain regions located within as well as outside the Neurologic Pain Signature (Fig. 2F). Nodes within the *h*^*2*^-component were spatially situated in the dorsal posterior insulae, anterior-, mid- and posterior cingulate cortex, precuneus, and orbitofrontal cortex.

### Genetic influence on behavioral sensitivity to electrical stimuli

There was a significant genetic influence on nociceptive thresholds, based on perception-matched aversive electrical stimuli (p<0.0001, *h*^*2*^ = 0.18, on choice of threshold). Estimates of the between-twin correlation of nociceptive thresholds for monozygotic twins was higher (*r* = 0.18, 95% CI = [-0.02-0.38]) than the between-twin correlation for dizygotic twins (*r* = 0.09, 95% CI = [-0.01-0.19]).

## Discussion

There is high variability in the way humans respond to nociceptive stimuli and express pain, yet there is little knowledge about the contributions of nature versus nurture to this variation. In this study, we used a twin-study approach to determine the magnitude and spatial representation of genetic influences on brain circuits involved in nociceptive processing. We found significant genetic influence on activity in brain regions typically activated by nociceptive processing (Fig. 1, Table 1). Interestingly, genetic influence on nociceptive functional connectivity was not restricted to these areas but also included regions across the brain (Fig. 2D-F).

**Table 1.** Genetic influence, *h*^*2*^ during nociceptive processing (*P<*0.05, family*-*wise error corrected). R = right hemisphere, L = left hemisphere.

| Area of Local Maximum | $h^2$ | MNI | | | Voxels in cluster |
| --- | --- | --- | --- | --- | --- |
|  |  | X | Y | Z |  |
| R Postcentral gyrus | .52 | 60 | -16 | 30 | 876 |
| R Insula | .50 | 36 | -20 | 18 |  |
| R Superior temporal gyrus | .45 | 54 | -30 | 18 |  |
| R Supramarginal gyrus | .44 | 56 | -24 | 18 |  |
| L Postcentral gyrus | .54 | -65 | -20 | 28 | 627 |
| L Supramarginal gyrus | .52 | -62 | -22 | 32 |  |
| L Insula | .43 | -36 | -20 | 18 |  |
| L Superior temporal gyrus | .43 | -64 | -30 | 22 |  |
| L Anterior cingulate | .46 | 0 | 20 | 28 | 620 |
| R Posterior-medial frontal | .41 | 2 | -14 | 58 |  |
| R Midcingulate | .40 | 8 | -14 | 40 |  |
| L Midcingulate | .40 | -2 | -4 | 40 |  |

Nociceptive responses in bilateral dorsal posterior insulae and mid/anterior cingulate cortex were influenced by genetics (Fig 1. Table 1), even if the cluster on the contralateral insular side was more pronounced. Previous studies have suggested that the dorsal posterior insulae may be of importance for nociceptive processing (Segerdahl, Mezue, Okell, Farrar, & Tracey, 2015). It is a primary projection point from the ventral medial nucleus of the thalamus and constitutes a core pathway for nociception in all primates (Craig, 2003). This thalamocortical pathway is believed to provide a sensory reflection of the condition of the body, and thereby has great evolutionary value (Craig, 2003). This is corroborated by fMRI data from newborn babies as it reveals a large overlap between nociceptive processing in adults and infants, including the thalamus, insulae and mid/anterior cingulate cortex (Goksan et al., 2015). This network could be considered as potential targets in studies searching for markers of chronic pain and novel treatment, especially for conditions with known familial risk. Genetic variability is likely to be involved in the mechanisms underlying some of our most common pain conditions (Parisien et al., 2017) but the mediating mechanisms are poorly understood. The results presented here demonstrate that nociceptive processing is significantly influenced by genetics and is likely to mediate the different nociceptive processing seen in individuals with chronic pain (Hashmi et al., 2013; Jensen et al., 2009).

Regarding the functional connectivity results, we observed that the nodes within the so-called *h*^*2*^-component (connectivity influenced by genetics) were localized in several different networks, most notably, the somatomotor, default-mode and dorsal attention networks (Fig. 2D-F). This indicates that genetic influence on functional connectivity during nociceptive processing encompasses both sensory and affective-cognitive processes. Since nociception is shaped by interactions between sensory, cognitive and affective processes there is indeed a possibility that some aspect of all these components is heritable. As an example, our results on the genetic influences on brain networks reveal a brain-wide pattern that includes regions implicated in cognitive-affective processes.

Notably, the largest number of connections in the *h*^*2*^-component was found between the default-mode and somatomotor networks (Fig. 2D). This was the case even though functional connectivity between the two was not the strongest (Fig. 2AC). In terms of functional connectivity, we observed a decrease in within default-mode network correlation, in line with previous findings (Kong et al., 2010; Kucyi et al., 2013). The clinical relevance for default-mode network has been observed previously since the precuneus region was associated with individual differences in pain sensitivity (Goffaux, Girard-Tremblay, Marchand, Daigle, & Whittingstall, 2014). Furthermore, functional connectivity was shown to decrease between default-mode network and primary somatosensory cortex following exacerbated pain in patients with chronic low back pain (Kim et al., 2019). We show an increase in functional connectivity between the default-mode and the somatomotor network, and that the largest number of connections were observed between the two in the heritable cluster. This is relevant to the translational potential between our data and clinical pain. The genetic influence on default-mode network and somatomotor connectivity, together with previous reports of altered connectivity in chronic pain, suggest it may serve as an intermediate marker of aberrant nociception.

Here, we isolated the genetic contribution to task-evoked functional connectivity. Yet, several findings show great similarity between task-evoked and resting-state functional connectivity (Cole, Bassett, Power, Braver, & Petersen, 2014; Fox & Raichle, 2007). Such similarities, however, should not be transferred by analogy to a comparison between resting-state and pain-evoked functional connectivity. Even comparing non-painful and painful stimuli shows marked differences whereby the former resembles a network formation akin to resting-state (Zheng et al., 2020). The cluster of edges identified in the present study captures variance associated with additive genetics supporting the search for a genetically informed neural pain signature (Davis et al., 2020). Future studies should compare resting-state and pain-evoked functional connectivity and estimate the extent of their shared genetics and the neural targets of their shared and non-shared genes.

As nociception is represented by activation in several brain regions, it has been difficult to determine which aspects of nociception are heritable and which ones are shaped by life experience. The data in the present study provides the first genetically informed nociceptive signature that distinguishes between heritable and acquired nociceptive responses in the brain. There is currently a need for better characterization of the biological and genetic foundations of the neural representation of pain. One major reason for the urgency of improving our understanding of the neural representations of pain is the opioid crisis, where opioid-based analgesics have created a wave of addiction, leading to overdoses and deaths. One review and a recent consensus paper by leading pain clinicians and scientists (Davis et al., 2020; Tracey, Woolf, & Andrews, 2019) explicitly ask for pain biomarkers– verifiable in preclinical models and patients. Stratification biomarkers may increase the probability of success in pharmacological clinical trials by as much as 21% in phase III clinical trials in all disease areas (Davis et al., 2020). Our results may help determine if clinical pain is manifested in genetically inferred nociceptive regions, and hopefully lead to beneficial sub-grouping and patient stratification.

There are several limitations in our study that need to be addressed. First, the experiment also included a fear conditioning task, which entails a risk that our findings are confounded by cognitive and affective processes related to learning and anxiety. On the one hand, our analytical approach isolated the effects of the nociceptive stimulus itself and hopefully minimized any brain activations related to the fear learning component of the experimental paradigm. On the other hand, there is an inherent affective component of nociception and it will thus be difficult to remove all fear-related brain activations as they may also be present during the nociceptive stimulation modeled in our analysis. Related to this, there are other psychological factors that are heritable, for example anxiety, that could influence nociceptive processing. We can, therefore, not exclude the contribution of closely related heritable factors to our findings. For example, genetic factors could influence anxiety that in turn influence nociceptive processing. Another limitation is that we examined the genetic influence on nociceptive processing and not subjective pain. While the nociceptive stimuli in our study represent aversive events in the sensory domain (Lee, Necka, & Atlas, 2020), participants did not provide subjective ratings of pain. This would have allowed a clearer relationship between genetic influences on brain activation and functional connectivity with the subjective pain experience. Further, this study examined only one nociceptive modality – electrical stimulation. Even if our findings elucidate heritable neural mechanisms that overlap with findings among patients with clinical pain they may not generalize to a clinical context. If we had used other nociceptive stimuli that stimulate deeper tissues, and provide C-fiber mediated activations, it would have made a stronger case for a possible clinical translation. Nevertheless, the level of the electrical stimuli in this study are comparable to previous studies that studied pain (Liu et al., 2020). Finally, the sample size is relatively small and may be underpowered to detect some effects. With our sample size, reaching 80% power (p_α_ = 0.05) requires the true effect of additive genetics to be 0.5. These calculations (Visscher, 2004; Visscher, Gordon, & Neale, 2008) are, however, based on a generic tool for twin studies and may not be comparable to the statistics of neuroimaging.

To summarize, our findings support the idea that brain regions associated with nociceptive processing are under significant genetic influence. The genetic influence on functional connectivity during nociceptive processing is not limited to core nociceptive brain regions, such as the dorsal posterior insulae and somatosensory areas, but also involves cognitive and affective brain circuitry. There are efforts to characterize the association between functional brain networks and gene expression (Richiardi et al., 2015). Future endeavors in the pain field can provide insights into clinical pain conditions and help improve their treatments.

## Author Contributions

J.R and F.Å designed the experiment. G.K and J.R performed the experiments. G.K and K.J wrote the first draft. All authors substantially revised the manuscript. X.C and T.N contributed with the software. G.K analyzed the data and G.K, X.C, W.H, T.N, I.T, P.F, F.Å and K.J contributed to interpretation of results. All authors have read and approved the manuscript.

### Data and materials availability

The data generated during the current study are available at osf.io with DOI 10.17605/OSF.IO/UWYEV. Further data and all the code used to produce figures and results are available at https://github.com/granitz/twin_pain.

## Supplementary Materials

Figures S1-S5

Tables S1

Captions for Data S1

