## supplementary_materials for "Genetic influences on brain activation and large-scale functional connectivity during nociceptive processing: a twin study"

**This PDF file includes:**

Figs. S1 to S5

Tables S1

Captions for Data S1

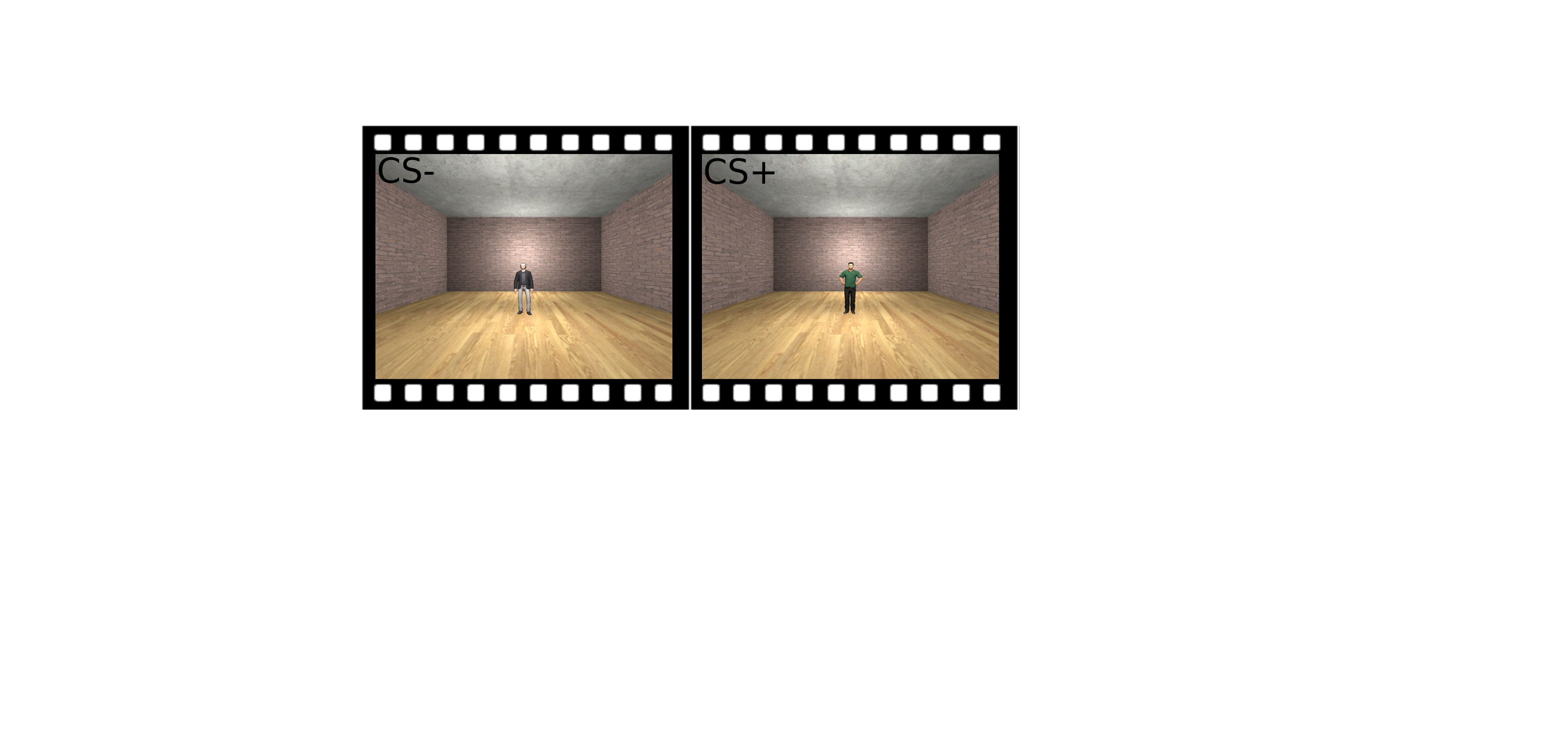

Fig. S1.

Experimental design. Two virtual characters served as conditioned stimuli (CS). One of the virtual characters served as aversive cue (CS+) and predicted the electrical shock and the other cue served as control (CS-). Each CS appeared for 6s with an inter-stimulus interval of 8-12s. Prior to the experiment, participants were not told which character would be associated with electrical shocks. During conditioning, each CS type was displayed 16 times. Eight of the CS+ presentations co-terminated with an electrical shock, and eight presentations of the same visual cue did not include a shock.

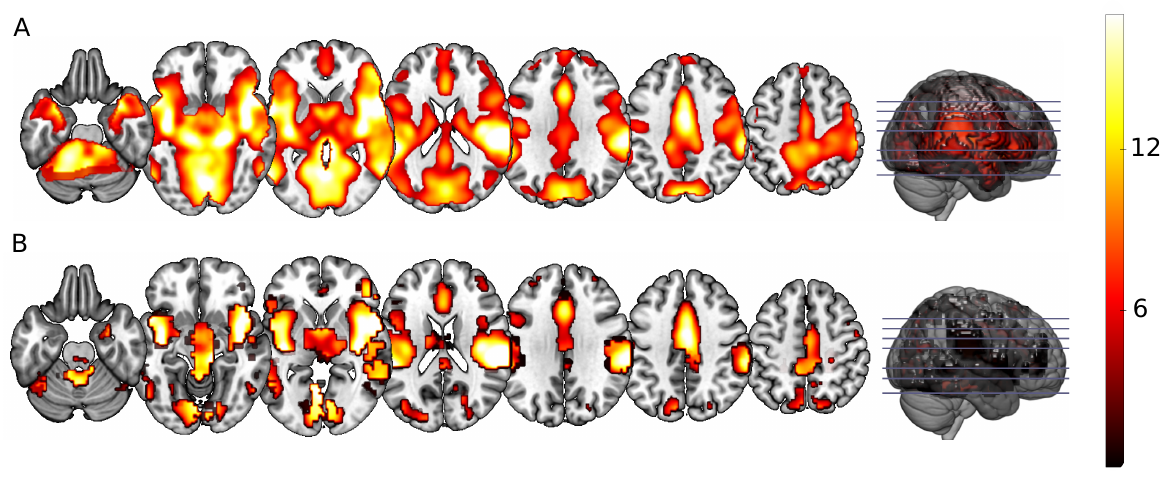
Fig. S2.

Brain responses during nociceptive processing. (A) Images represent whole-brain activity to nociceptive stimuli (p<0.05, *FWE* corrected). (B) Same as in (A) but here limited to regions within the borders defined by the Neurologic Pain Signature.

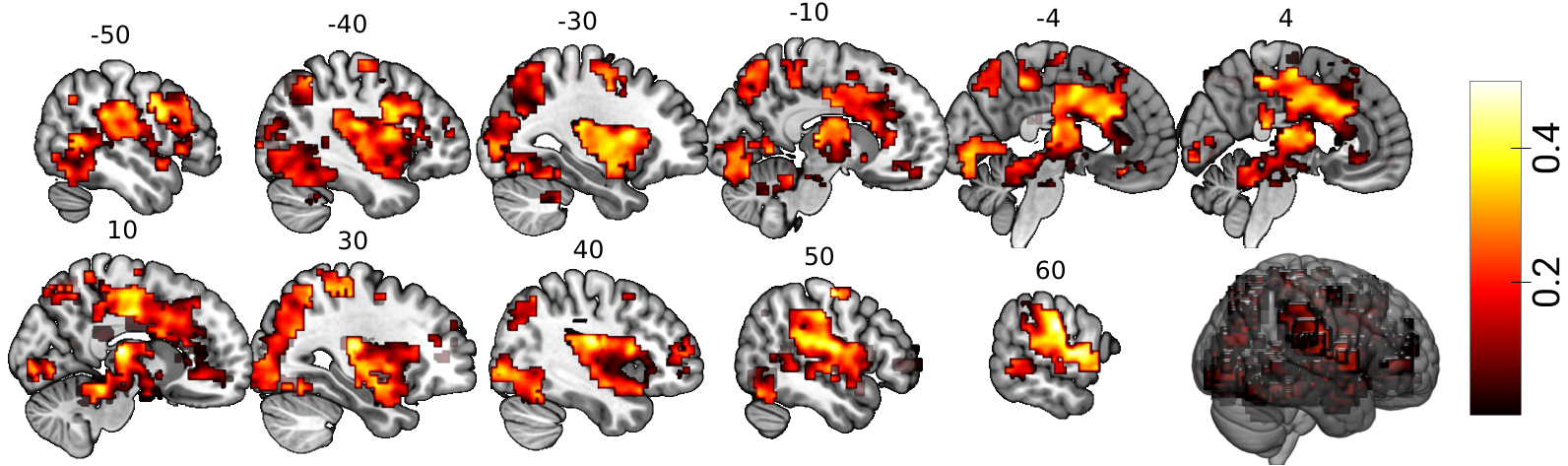

Fig. S3.

Unthresholded genetic influence, *h^2^*. Genetic influence was estimated per voxel on individual level contrast images masked with the Neurologic Pain Signature.

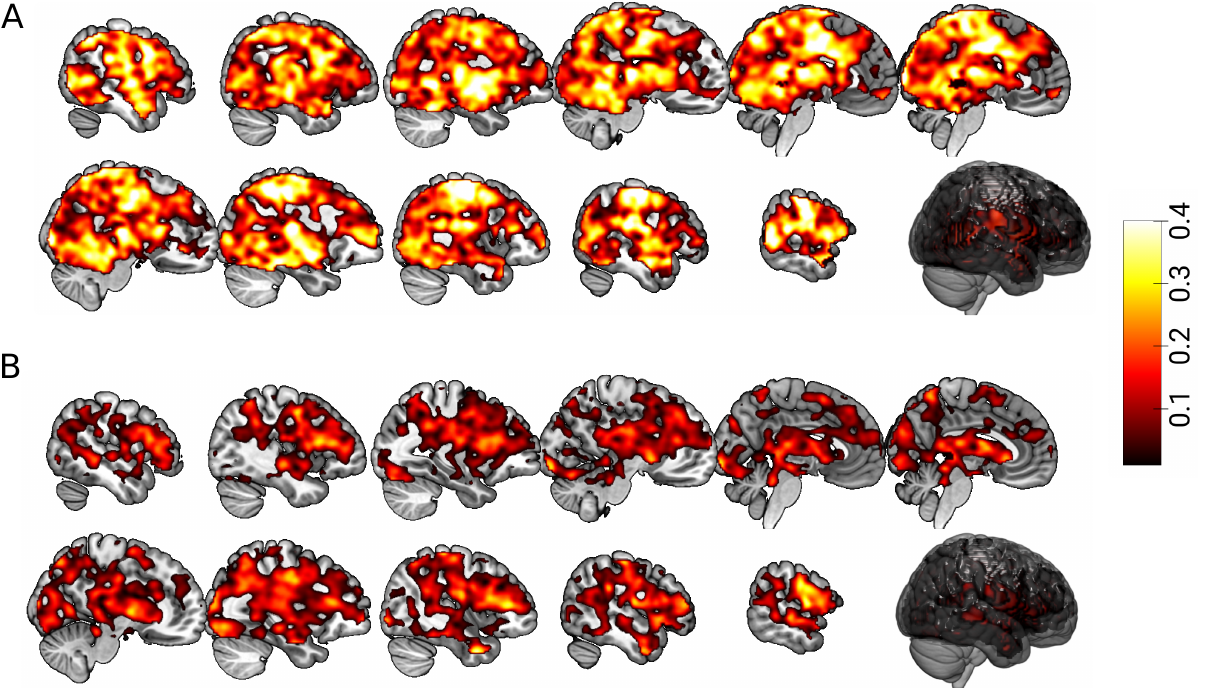

Fig. S4.

Unthresholded whole-brain between twin-pair correlations for (A) identical or monozygotic and (B) fraternal or dizygotic twin-pairs.

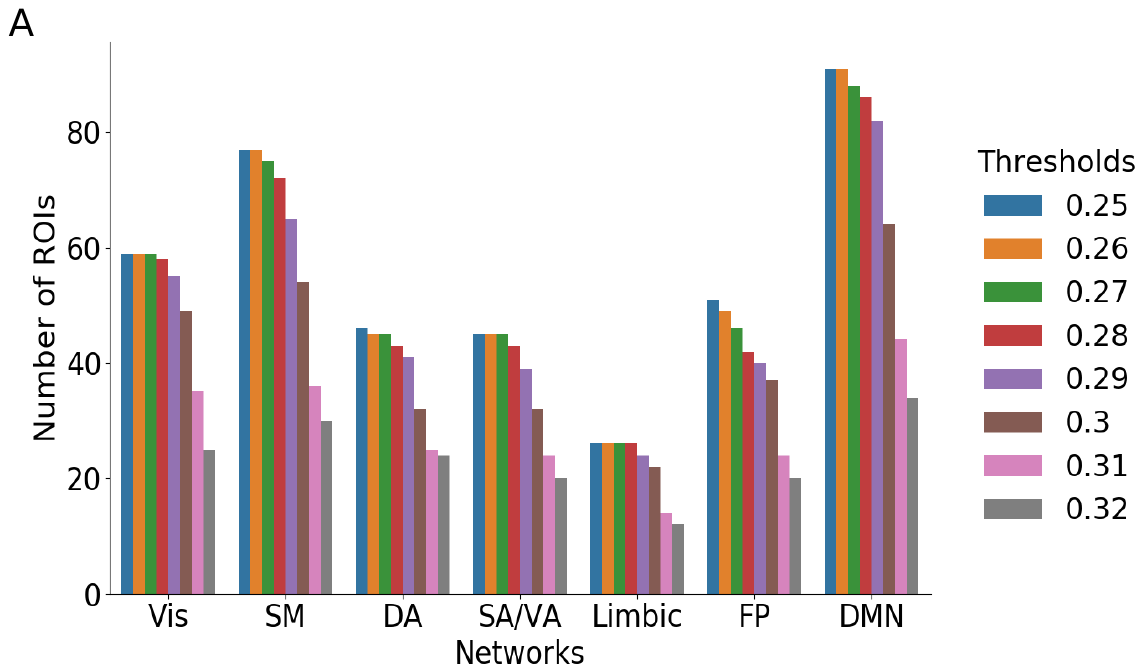

Fig. S5.

The *h^2^*-components for different thresholds of *h^2^*. With a lower threshold value, larger but weaker connected components are identified.

Table S1.

Brain areas with greater fMRI response during nociceptive processing (*P <0.05, family-wise error corrected*). R = right hemisphere, L = left hemisphere. Note that SPM Anatomy identified only one cluster.

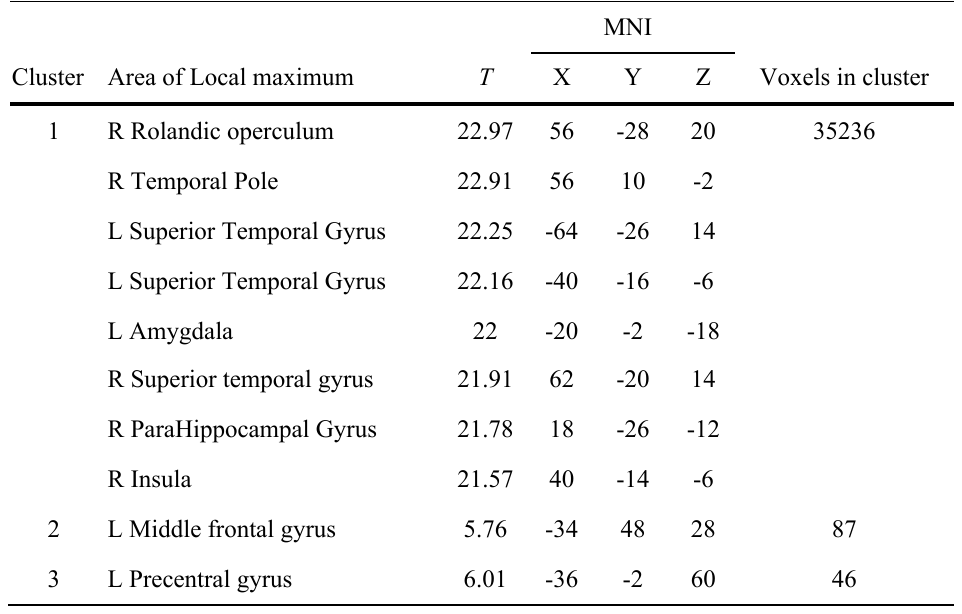

Data S1. (separate file)

Niftii files corresponding to unthresholded a^2^, c^2^ and e^2^ that are pictured in *SI Appendix,* Fig S3 are available at https://github.com/granitz/twin_pain.
